# Disulfide engineering reveals unexpected pro- and anti-aggregation conformers of human α-synuclein

**DOI:** 10.64898/2026.01.07.698305

**Authors:** Aslam Uddin, Eugene Serebryany

## Abstract

Intrinsically disordered proteins can aggregate in many distinct conformations (polymorphs). Polymorphs are a striking example of fold-switching: one primary structure able to form distinct tertiary structures. Distinct polymorphs can yield distinct molecular, cellular, and disease phenotypes. Disulfide crosslinks canalize disordered proteins into distinct regions of the conformational landscape, even if the monomer remains disordered. Human α-synuclein is a disordered, natively Cys-free protein involved in synaptic transmission. Its amyloid aggregation is implicated in Parkinson’s disease and other synucleinopathies. Recent cryo-EM fibril structures have revealed distinct classes of amyloid polymorphs that wild-type human α-synuclein can adopt. We created three double-Cys α-synuclein variants predicted to form intramolecular disulfide bridges compatible with known amyloid polymorphs, plus three other double-Cys variants at arbitrary positions as controls. We purified all six variants as intramolecularly disulfide-crosslinked monomers in solution. Two of the three predicted pro-aggregation variants formed amyloid fibrils as expected, and two of the three control variants did not. Surprisingly, one of the controls formed amyloid fibrils with a distinctive morphology by negative-stain TEM, suggesting the possibility of a previously unknown polymorph. Conversely, the crosslinked variant that failed to aggregate as expected showed sub-stoichiometric, dose-dependent anti-aggregation activity, strongly suppressing amyloid formation by wild-type α-synuclein.

## Introduction

Human α-synuclein is a 140 residue intrinsically disordered protein (IDP), known to adopt distinct conformations in the amyloid aggregated state [1–4]. It is natively abundant in presynaptic terminals of neurons and regulates diverse neuronal functions, such as synaptic transmission and plasticity [5–7]. However, some of the conformations α-synuclein can adopt result in the formation of aggregates or toxic oligomers characteristic of Parkinson’s disease and other neurodegenerative disorders, such as dementia with Lewy bodies and multiple system atrophy (MSA) [6, 8, 9]. Experimental elucidation of the properties of α-synuclein’s many possible conformations is challenging due to a dearth of tools for studying disordered conformations [10–12]. Yet, it is critically important for understanding the conformational basis of the associated neurological diseases.

Disulfide crosslinks in an IDP can dramatically alter their molecular and cellular phenotypes by constraining the region of conformational space accessible to it, even if it remains disordered. Thus, Serebryany *et al.* have shown that intramolecular disulfide trapping of HdeA, an IDP in the *E. coli* periplasm, can shift both the protein’s molecular phenotype (e.g., hydrophobicity) and the phenotype of the bacterium itself (e.g., the likelihood of cell lysis) [13]. Walker *et al.* used disulfide trapping to probe structural features that affect the aggregation propensity of tau [14]. Similarly, Carija *et al.* demonstrated that introducing a single disulfide into the natively Cys-free wild-type human α-synuclein resulted in a more compact conformation and altered aggregation behavior [15]. We set out to use intramolecular disulfide crosslinking more systematically to constrain the natively disordered conformational ensemble of α-synuclein to favor or disfavor aggregation.

Here, we took advantage of cryo-EM structures of α-synuclein fibrils to design three double-Cys variants where disulfide formation was expected to promote amyloid aggregation and compared them three randomly chosen double-Cys variants. The sequence of α-synuclein is conventionally divided into three domains. The N-terminal domain (residue 1 to 60) promotes binding to synaptic vesicles [15], and the loss of this function due to α-synuclein aggregation is thought to contribute to neuropathology [16]. The NAC (non-amyloid-β component) domain comprises residues 61-95, is highly hydrophobic, and drives α-synuclein aggregation [17]. The C-terminal domain is highly charged, so it typically retains mobility in α-synuclein oligomers [18, 19] and fibrils [20, 21]; it may also bind synaptic vesicles via bridging Ca²□ions [22]. Nonetheless, cryo-EM structures reveal that the amyloid fibril cores consist not only of the NAC domain, but also significant parts of the N-terminal domain and even parts of the C-terminal domain [23]. We therefore chose the six variants (see Experimental Design) without limiting our search to the NAC domain alone.

We quantified the aggregation behaviors of these six variants and their interactions with the wild-type protein. We used kinetic models to distinguish among effects of disulfide trapping on primary nucleation, secondary nucleation, and elongation. We used transmission electron microscopy (TEM) to analyze how intramolecular disulfide trapping changed overall aggregate morphology. We found that two of the three variants that were expected to aggregate did so, while two of the three randomly chosen variants had undetectable aggregation. However, aggregation-resistant disulfide-trapped α-synuclein variants also inhibited amyloid aggregation of the WT α-synuclein. Surprisingly, the strongest anti-aggregation effect was from the variant that was itself expected to aggregate but did not. Meanwhile, the variant that had been expected not to aggregate but did, albeit slowly, showed a distinctive aggregate morphology, and we verified that its disulfide is not consistent with any known amyloid polymorph of α-synuclein.

Our results highlight how conformational constraints in IDPs can alter both their supramolecular behavior and that of the otherwise unconstrained WT. They also highlight that we are still far from having fully explored the conformational landscapes of IDPs in terms of their molecular and supramolecular phenotypes. While disulfides can be labile *in vivo*, other chemical crosslinks are more stable. Disulfide engineering can guide other forms of covalent crosslink engineering. Our results suggest that conformationally constrained versions of human α-synuclein could potentially be used to modify the progression of sporadic synucleinopathies.

## Results

### Experimental design

To elucidate the mechanisms underlying the aggregation of conformationally constrained α-synuclein, we engineered six disulfide-crosslinked variants by introducing mutations to Cys within the protein sequence spanning residues 22 to 95. Of the six engineered double-cysteine variants, three (T22C-T33C, T44C-T54C and T64C-T81C) were designed with cysteine residues arbitrarily positioned within the sequence spanning residues 22–95 to assess the impact of non-specific placement on α-synuclein aggregation (**Figure 2a, b).** The remaining three variants (A29C-V71C, A69C-A91C and E46C-K80C) were guided by structural insights derived from existing cryo-EM polymorphs (PDB IDs: 6CU7, 8PK4), enabling a comparative analysis between structurally informed and randomly positioned mutations (**Figure 2a, b**). For simplicity, we will refer to these variants by their Cys positions: **22-33**, **44-54**, **64-81**, **29-71**, **69-91**, and **46-80**.

### Characterization of oxidized α-synuclein double-Cys variants

Prior to initiating aggregation assays, we oxidized the double-cysteine variants by incubating them overnight at 4□°C in 20□mM phosphate buffer (pH 7.0) containing 1□mM oxidized glutathione (GSSG). To remove glutathione and prevent future thiol-disulfide exchange or glutathionylation, the samples were subsequently purified using size exclusion chromatography (**Figure 1a**). To verify that oxidation of double-cysteine α-synuclein variants was primarily intramolecular, maleimide-PEG labeling combined with SDS-PAGE was used (**Figure 1b**). Maleimide reacts with free thiol groups, and PEGylation introduces a detectable gel shift. This shift allows estimation of the number of free thiols per protein molecule [24]. Proteins with no shift are fully oxidized (intramolecular disulfides), while shifted bands indicate remaining free thiols (**Figure 1c**). This semi-quantitative method confirmed that the oxidation strategy yielded intramolecular disulfide bonds, preserving monomeric structure while introducing conformational constraints. Note that any intermolecular disulfides would have resulted in covalent dimers of the proteins, which would have been fully removed during the size exclusion chromatography step.

**Figure 1:**
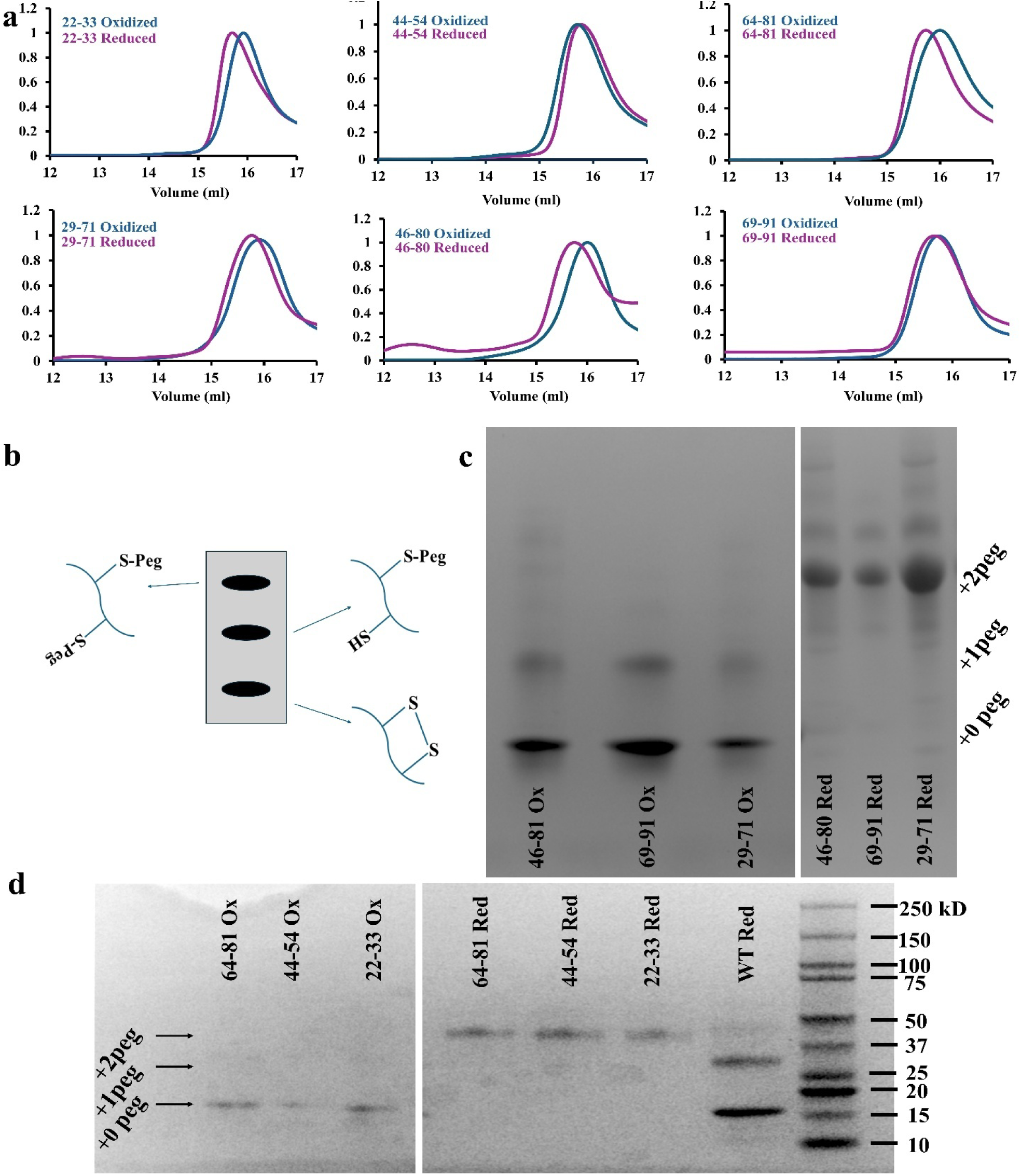
Characterization of oxidized and reduced α-synuclein double-Cys variants. **(a)** Size exclusion chromatography traces of oxidized (disulfide-trapped) vs. reduced variants showed a consistent slight shift to later elution volumes for five of the six variants. This shift is expected for a monomeric IDP with a more compact conformational ensemble. **(b)** Schematic representation of the PEGylation gel shift assay for quantifying the number of free thiol groups per protein molecule. This method has an advantage over Ellman’s test because it measures the distributions of free and disulfide-bonded Cys across the molecular population, rather than only the average [24]. **(c)** and **(d)** SDS-PAGE gels of the six variant proteins and the WT as control, incubated with maleimide-PEG to quantify the distribution of the number of free thiol groups per protein molecule. Note that the WT lacks thiol groups (it is Cys-free), so the minor bands observed at the +1PEG and +2 PEG levels in WT in **(d)** are an artifact of the maleimide reacting with a non-thiol moiety on the protein. They are nonetheless useful as standards.

**Figure 2:**
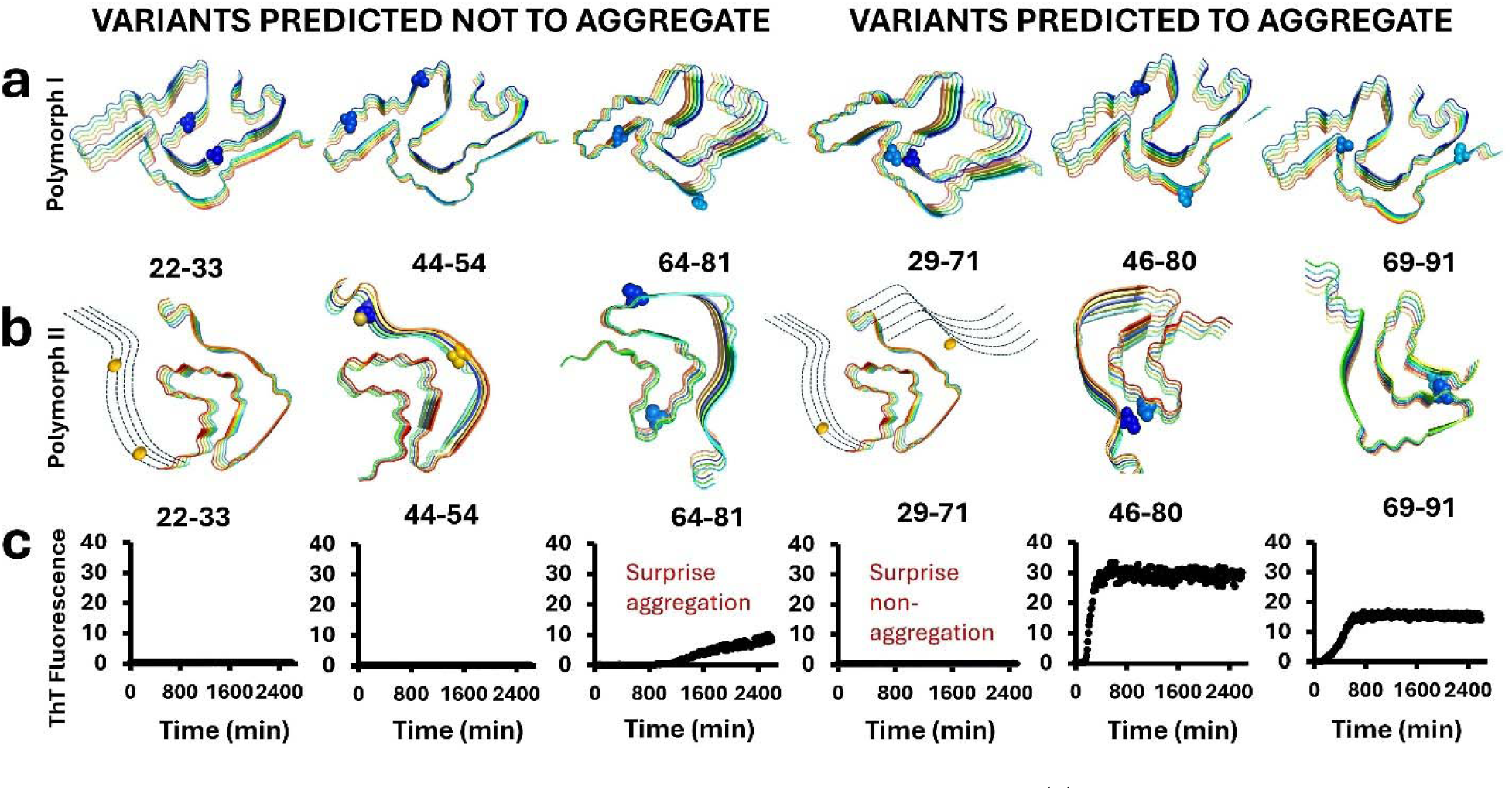
Disulfide engineering of human α-synuclein aggregation. **(a)** Mutation sites in six double-Cys variants shown on a representative structure of one major polymorph class (polymorph I, PDB ID: 8PK4). **(b)** The same on the other major polymorph class (polymorph II, PDB ID: 6CU7). Some mutation sites were outside the structured core of this polymorph, so we depicted their locations schematically (*dashed lines*). **(c)** Experimental amyloid aggregation propensities (measured by thioflavin T fluorescence) for the six double-Cys variants.

To further confirm the redox state of the variants, we used electrospray ionization mass spectrometry (ESI-MS) to measure their exact masses. The measured molecular masses were in excellent agreement with the theoretical ones. Observed molecular weights (**Table S1** and **Figure S1**) for all variants matched the predicted mass of the oxidized form of the respective proteins, modulo a calibration error of -0.5 or -0.6 Da. in all cases. The WT protein, which has no disulfides, matched its theoretical molecular weight with the same shift of -0.5 Da. as the oxidized variants. This indicates our oxidation procedure resulted in ∼100% disulfide bonding in the variants. However, we observed a second major peak at +30 Da. for variant **46-80**. Considering the presence of additional minor peaks in that sample, including those consistent with a phosphate or sulfate adducts (**Figure S1**), this +30 Da. peak in variant 46-80 may have resulted from on-column contaminants or incomplete desalting during ESI-MS analysis.

### Determination of variant aggregation propensities and aggregate morphologies

**Figure 2a,b** shows the locations of the introduced Cys residues in the six double-mutants we designed. The locations are depicted on representative structures of the two major classes of α-synuclein amyloid polymorphs. We measured amyloid aggregation by monitoring the increase in thioflavin T (ThT) fluorescence intensity over time (**Figure 2c**). Most results aligned with our structure-guided expectations, but not all. Two of the three randomly designed mutants (**22-33** and **44-54**) did not aggregate, but the third (**64-81**) did, albeit slowly. This variant’s aggregates were also morphologically distinct from the others (**Figure 3**). Conversely, aggregation was observed in two of the three mutants designed to aggregate based on existing Cryo-EM polymorphs, but the third (**29-71**) did not aggregate (**Figure 2c**).

**Figure 3:**
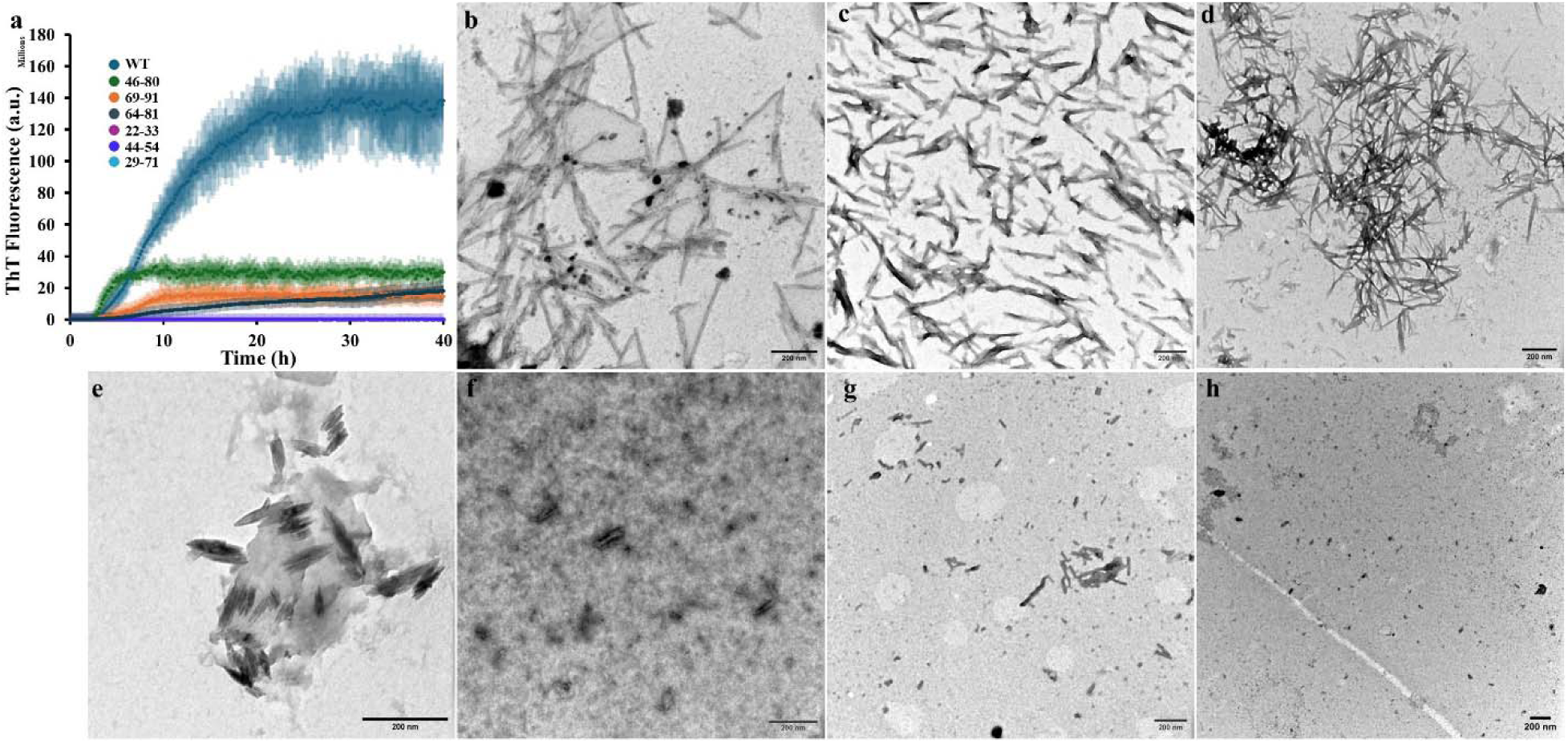
**(a)** Aggregation kinetics of α-synuclein WT, **64-81**, **69-91** and **46-80** assessed by ThT fluorescence over time. All protein concentrations were 80 μM. Transmission electron micrographs (TEM) showed that α-synuclein variants formed morphologically distinct aggregates: (**b**) WT α-syn (**c**) **22-33** (**d**) **44-54** (**e**) **29-71** (**f**) **64-81** (**g**) **69-91** (**h**) **46-80**. Scale bars in all TEM images represent 100 nm.

While variants **22-33** and **44-54** exhibited no aggregation in the oxidized state (**Figure 2c**), they did aggregate in the presence of reducing agent (**Figure S2**). Hence, their aggregation resistance was attributable to the conformational constraint of their disulfide bonds and not merely to altered protein sequence. Variants **46-80**, **64-81**, and **69-91** introduced changes to the non-amyloid-*β* component (NAC) region, which spans residues 61–95. This region is known to be crucial for aggregation into amyloid fibrils [25], and some mutations within or near it are known to affect aggregation [26]. In variant 46-80, however, the newly introduced disulfide replaces a salt bridge between E46 and K80 that exists within the known polymorph II class (**Figure 2a**). Accordingly, this variant aggregated even more rapidly than WT, though the maximum amplitude of ThT fluorescence was below that of the WT for all variants (**Figure 3a**).

TEM analysis (**Figure 3b-h**) revealed that the three α-Synuclein mutants (**22-33**, **44-54** and **29-71**) which showed no aggregation in ThT assays also lacked visible aggregates, confirming their non-aggregating behavior. In contrast, the mutants (**46-80**, **69-91** and **64-81**) with aggregation propensity formed morphologically distinct aggregates compared to wild type, suggesting that sequence-specific variations influence not only aggregation kinetics but also aggregate structure.

The **64-81** variant’s behavior is particularly interesting, since it formed amyloid aggregates despite not being predicted to do so based on the available major classes of amyloid structures. Therefore, we systematically investigated whether the structure of the crosslinked **64–81** variant corresponds to any of the existing cryo-EM polymorphic structures of α-synuclein (**Figure 4**). The C_β_-C_β_ distance characteristic of a disulfide bridge is approximately 5 Å. Two recent studies classify all currently known α-synuclein amyloid polymorphs (>100 distinct cryo-EM structures) into several classes [27, 28]. We systematically measured the C_β_-C_β_ distance between residues 64 and 81 for representative structures from all the known classes of α-synuclein amyloid polymorphs, as well as from all currently unclassified polymorphs (**Figure 4**). Strikingly, none of these polymorph structures were consistent with aligned with a 64–81 disulfide. suggesting that this crosslinked variant may adopt a previously uncharacterized α-synuclein polymorph.

**Figure 4:**
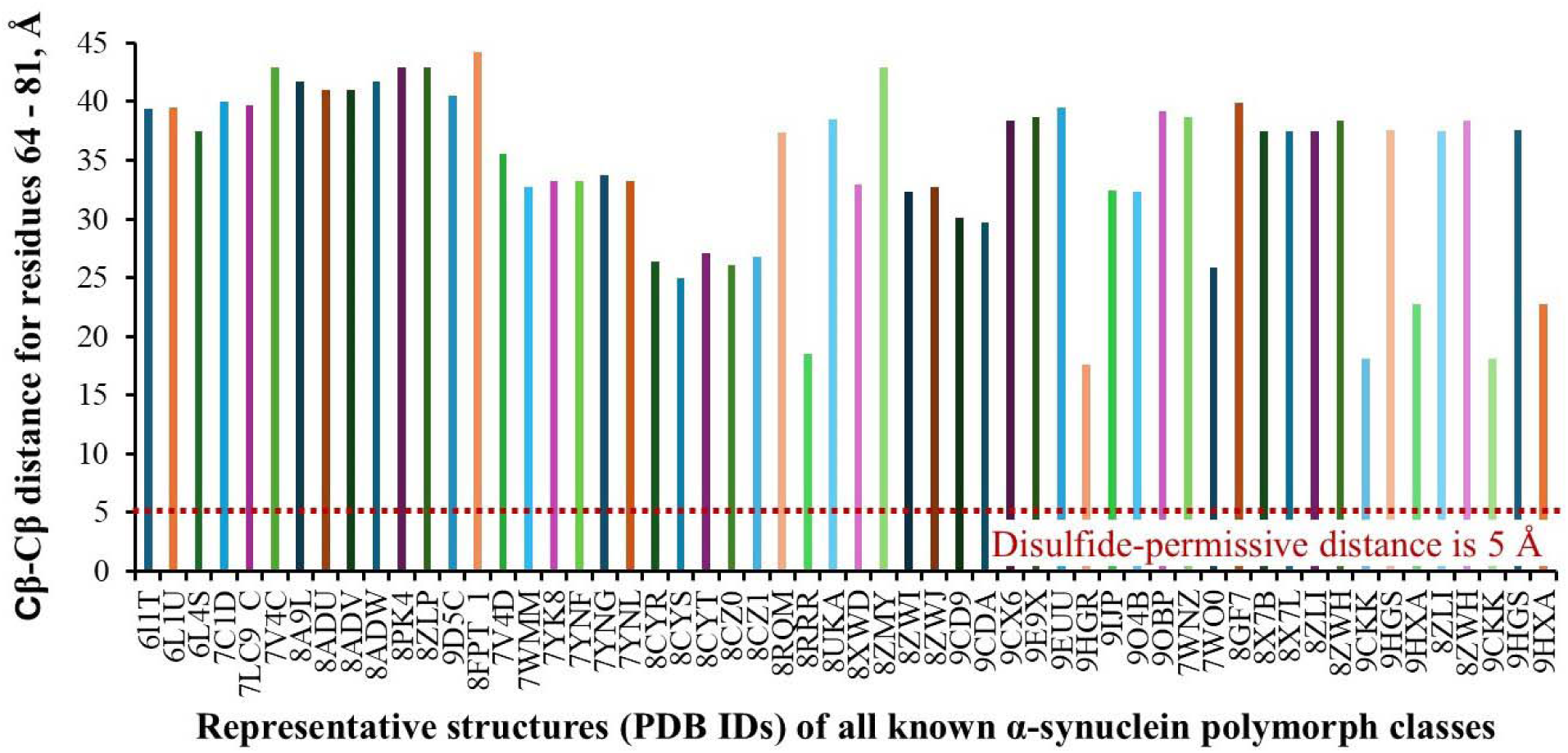
Variant 64-81 may represent a previously unknown amyloid polymorph of α-synuclein. The measured distances between the β-Carbon atoms of residues 64 and 81 in representatives of all currently known polymorph classes, along with all currently unclassified polymorphs, were much larger than the disulfide-permissive distance.

The aggregation kinetics of α-synuclein variants that exhibited fibril formation were quantitatively analyzed using the **AmyloFit** platform [29], a web-based tool for global fitting of protein aggregation data to mechanistic models (**Figure 5**). This analytical approach establishes a direct link between macroscopic observations of protein aggregation and the fundamental microscopic mechanisms. This connection facilitates the evaluation of the nucleation and growth processes and enables the determination of their corresponding rate constants. To achieve optimal global fitting, we evaluated scaling exponents and applied kinetic models incorporating saturation elongation and fragmentation mechanisms. This approach enabled the extraction of microscopic rate constants ***k_n_k*_+_** and ***k_+_k*_-_** that characterize the underlying aggregation pathways. In all cases, the reaction order of primary nucleation (nc) was fixed at 2.

**Figure 5:**
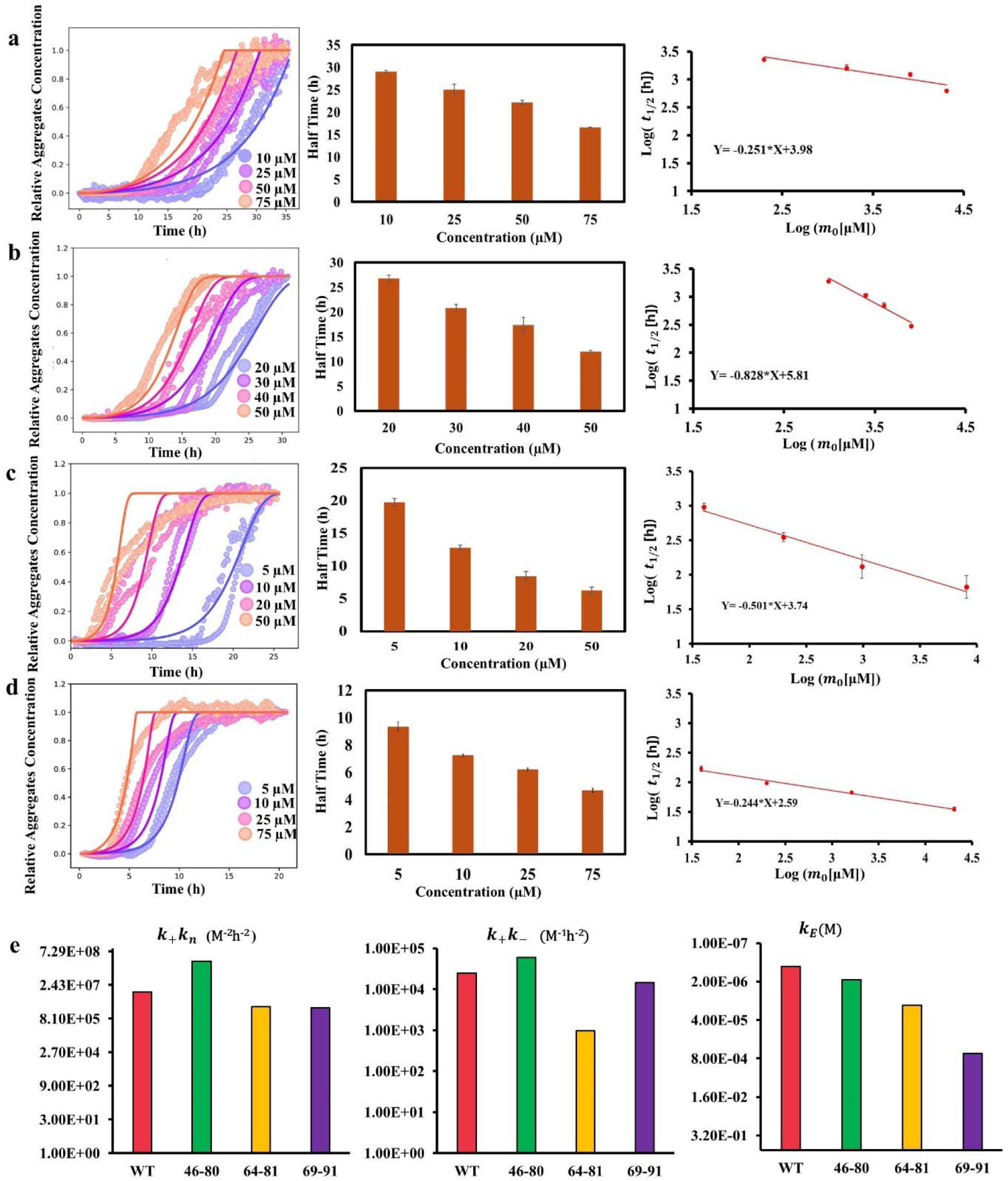
Aggregation kinetics of α-synuclein variants WT, 64-81, 69-91 and 46-80 assessed by ThT fluorescence over time. (**a, b, c, d**) Normalized traces were fitted using AmyloFit (*Left panel*) with the Saturating Elongation and Fragmentation model [29] (*solid lines*). The aggregation half-times (t_1/2_) exhibits a clear concentration dependence (*Central panels*), and double-logarithmic plots (*Right panels*) were used to determine the scaling exponent (i.e., the slope of linear regression function) for selection of the appropriate aggregation model. **e)** Kinetic parameters derived from the fitting are ***k_n_k*_+_** (primary nucleation and elongation), ***k_+_k*_-_**(elongation and fragmentation), ***K_e_*** (concentration of half-maximal elongation speed).

The combined rate constant for primary nucleation and fibril elongation (*k_n_k*_+_) was markedly higher in the **46–80** variant (2.75 × 10□) compared to wild-type α-synuclein (1.19 × 10□) (**Figure 5e**). In contrast, the **69–91** and **64–81** variants exhibited substantially lower values (2.50 × 10 and 2.75 × 10□, respectively), nearly an order of magnitude below wild type. Similarly, the combined contributions of fibril elongation and fragmentation (*k_+_k*_-_) were elevated in the **46–80** variant (5.93 × 10□) relative to wild type (2.5 × 10□), while the 69–81 mutant (1.47 × 10□) was nearly equivalent to wild type, and the **64–81** variant (9.59 × 10²) was significantly lower, further supporting the differential aggregation behavior among the mutants. Overall, we observed that intramolecular disulfide bonds constraining the protein’s conformation may inhibit or alter aggregation pathways and modulate nucleation and elongation kinetics compared to the wild-type protein. In short, disulfides can control not only aggregate structure, but also the kinetics.

### Effect of non-aggregating mutants on the aggregation of wild type α-synuclein

Carija *et al.* reported that a disulfide-trapped α-synuclein variant had an anti-aggregation effect [15]. Therefore, we evaluated the influence of non-aggregating mutants (**22–33**, **44–54**, and **29–71**) on the aggregation kinetics of wild-type α-synuclein. We conducted a series of aggregation assays in which wild-type α-synuclein was incubated in the presence and absence of increasing concentrations of the variants. Aggregation reactions were performed as above, in 96-well plates with Teflon balls to provide mechanical agitation, thereby enabling continuous and parallel monitoring of Thioflavin-T (ThT) fluorescence emission across all samples. The resulting kinetic profiles (**Figure 6a-c**) revealed that two of the non-aggregating mutants **44-54** and **29-71** exerted a pronounced modulatory effect on WT α-synuclein aggregation with concentration dependent manner, whereas one mutant did not significantly alter the aggregation behavior of the wt. protein.

**Figure 6:**
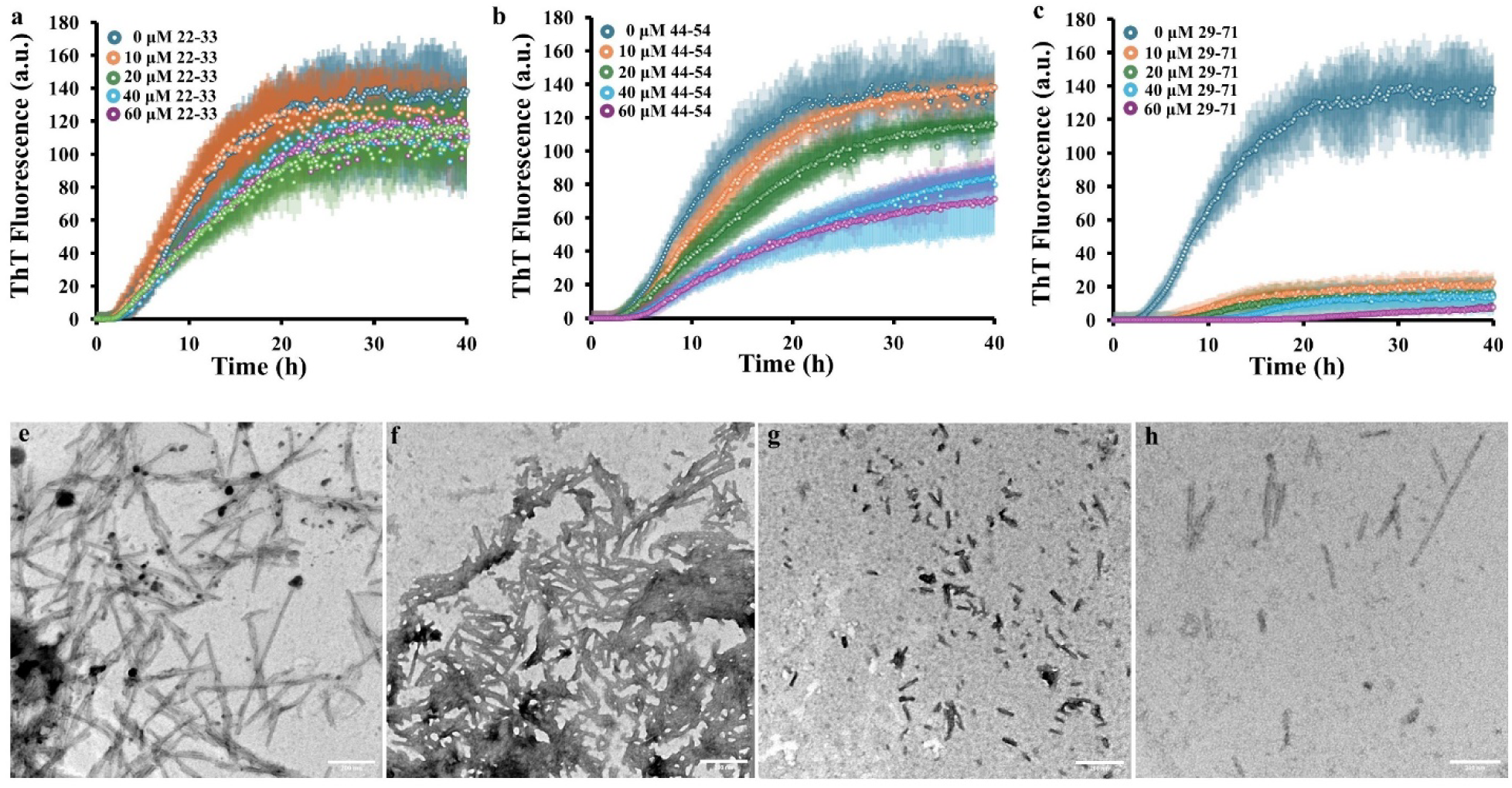
Aggregation kinetics of WT α-synuclein in the presence and absence of non-aggregating mutants: **(a) 22–33** and **(b) 44–54**, **(c) 29-71**, **(d-g)** TEM images of WT α-Syn in the absence **(d)** and presence of **e) 22-33**, **f) 44-54**, **g) 29-71**.

Transmission electron microscopy (TEM) revealed fewer amyloid fibrils in WT α-synuclein samples incubated with **29-71** or **44–54** (**Figure 6f, g**), indicating strong inhibition of aggregation. By contrast, variant **22-33** (**Figure 6h**) did not noticeably affect fibril density, which is consistent with its much more modest effect on the WT’s aggregation kinetics.

The aggregation kinetics of WT *α*-synuclein, with and without non-aggregating variants, were analyzed quantitatively by fitting a multistep secondary nucleation model using the **AmyloFit** platform [29]. This approach enabled the extraction of the microscopic rate constant (*knk*+) and (*k_2_k*_+_), referred to as the combined rate constants for the primary and secondary pathways, respectively, and *K_M_* is the Michaelis-Menten constant (**Figure 7**). Quantitative analyses of the fitted data revealed that the non-aggregating mutants **29-71** specifically modulate the primary nucleation pathway (*knk*+) during wild-type α-synuclein aggregation, whereas mutant **44-54** predominantly inhibits secondary nucleation. Notably, mutant **22-33** demonstrated no significant effect on any of the aggregation pathways evaluated.

**Figure 7.**
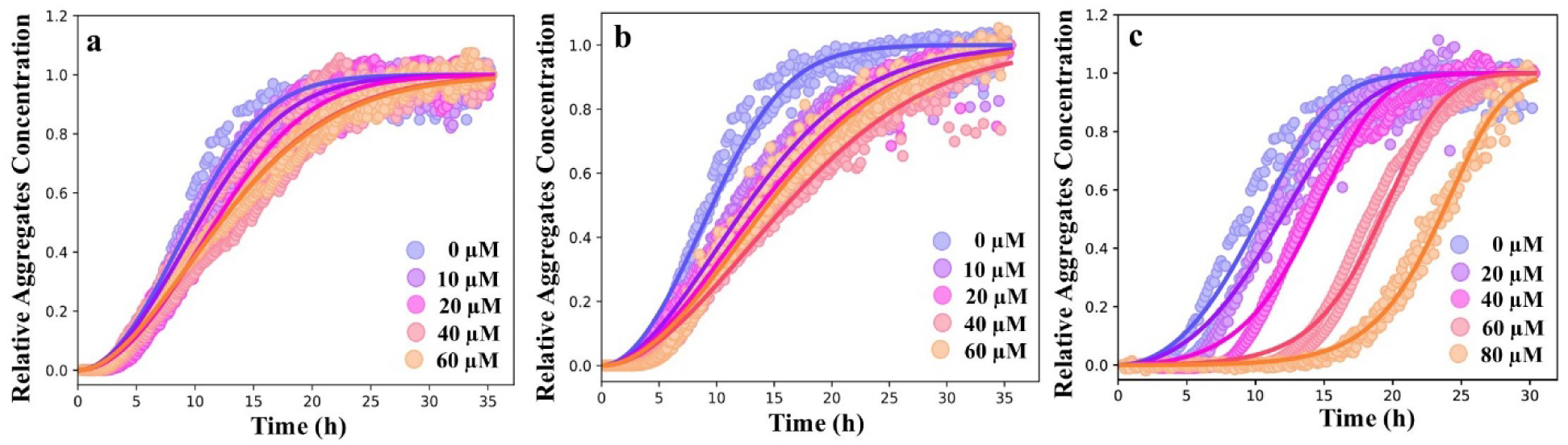
Non-aggregating variants have distinct inhibitory effects on WT α-synuclein aggregation. Aggregation kinetics were normalized and fitted with **AmyloFit** using the multistep secondary nucleation model (*solid lines*) in the presence or absence of non-aggregating variants: (**a**) **22-33**, (**b**) **44-54**, and (**c**) **29-71**.

## Discussion

The aggregation of α-synuclein into amyloid fibrils is a key factor in Parkinson’s disease and other synucleinopathies [30, 31]. Distinct amyloid polymorphs are known to cause distinct phenotypes in synucleinopathies [32], tauopathies [33, 34], and other neurodegenerative diseases [35, 36]. It is therefore valuable to characterize as many polymorphs of α-synuclein as possible. This presents a challenge because α-synuclein is an intrinsically disordered protein (IDP), so, in principle, almost any conformation is possible. Recent advancements in cryo-electron microscopy (Cryo-EM) have allowed researchers to identify two major classes of α-synuclein polymorphs, but also a variety of smaller polymorph classes [27, 28].

Disulfide engineering is a well-established approach for stabilizing protein conformations [37]. Conversely, disulfide scanning has been used to probe protein aggregation pathways, e.g., of the prion protein [38]. More recently, high-throughput disulfide scanning has been developed to map the conformational landscape of an intrinsically disorder protein to its phenotypic landscape [13]. Naturally occurring disulfides have revealed key insights into the aggregation pathways of many proteins, such as lens γ-crystallins [39–41], superoxide dismutase [42–44], and tau [14, 45, 46]. Disulfide engineering of α-synuclein could be a very fruitful strategy for constraining the protein to explore distinct conformers and measuring their aggregation. However, this has not been attempted in any systematic way: we are aware of only one previous study reporting one disulfide-bridged α-synuclein variant [15] and one reporting a double-disulfide variant [47]. The main reason is likely that structures were not available until recently to guide disulfide engineering studies. Now that cryo-EM structures of the major amyloid polymorphs are known and structural classifications are available, we have carried out this disulfide-trapping experiment to explore the link between conformation within the polymorph and aggregation propensity.

Introducing the mutations to Cys could, in principle, alter the conformational landscape on its own. The double-Cys variants that did not aggregate when oxidized did aggregate under reducing conditions (**Figure S2**). Future studies should examine whether variants like **64-81** can seed WT α-synuclein aggregation at least under some conditions. If so, then the natively Cys-free WT may adopt the same novel conformer as the disulfide-constrained variant. However, a study of the effect of naturally occurring intramolecular disulfides on tau aggregation revealed significant seeding barriers between oxidized and reduced conformations [45].

For a properly controlled study, we designed six double-cysteine variants. Three of the variants (**46-80**, **69-91** and **29-71**) were predicted to form intramolecular disulfides within one of the two known major amyloid polymorphs, while the other three variants (**22-33, 44-54** and **64-81**) were chosen arbitrarily as controls. The residue pairs in the control set do not approach each other in the known tertiary structure, precluding intramolecular disulfides linking them in the aggregated state. However, the soluble monomer is disordered, so we expected any pair of Cys residues to readily form an intramolecular disulfide bond in dilute protein samples, and this was, indeed, what we observed (**Figure 1**).

Four of the six disulfide variants behaved qualitatively as predicted: two from the first group and two from the second. The kinetic analysis demonstrated that the formation of disulfide bonds between distant regions of the polypeptide chain significantly alters the microscopic rate constants associated with amyloid fibril formation. These findings suggest that covalent linkages introduced through disulfide bonds can modulate the conformational dynamics of distinct α-synuclein variants, thereby influencing the rate of microscopic step involved in the pathway of fibril assembly.

However, the remaining two variants showed unexpected behavior. The inability of one designed variant to aggregate, despite its predicted aggregation-competence, suggests that subtle structural constraints or local sequence context may significantly influence amyloid formation. What was more surprising is that this variant (**29-71**) not only avoided aggregation itself but also strongly inhibited the aggregation of WT α-synuclein in a concentration dependent manner (**Figure 6c**). The non-aggregating variant **44-54** also acted as an aggregation suppressor of the WT, whereas **22-33** had only minimal impact (**Figure 6b, a**). Only one prior study has reported a modest chaperone-like effect from a disulfide-trapped α-synuclein variant [15]. The effects observed in the present study for variants **29-71** and **44-54** are substantially larger and suggest that this chaperone-like effect of constrained α-synuclein conformers on unconstrained ones may be widespread across the conformational landscape.

Moreover, the aggregation suppression effects were distinct: **44-54** primarily reduced the growth rate of WT aggregation, but **29-71** primarily increased the lag time (**Figure 7**). The latter variant also strongly suppressed the maximum aggregation amplitude that could be measured on the experimental time scale (**Figure 6c**). Both aggregation suppression effects occurred at sub-stoichiometric ratios of the suppressor variant to the WT. The observation that conformationally constrained variants can suppress distinct aspects of WT aggregation depending on the location of the constraint raises two intriguing possibilities. First, disulfide scanning could be a unique way to dissect the kinetic mechanism of aggregation even without modifying the WT protein in any way. Second, a cocktail of distinctly constrained variants could act as a potent all-around suppressor aggregation of the WT protein’s aggregation.

The mechanisms of aggregation suppression remain to be fully elucidated in future studies. We hypothesize that the 44-54 and 29-71 disulfides constrain the protein to conformers sufficiently similar to a WT aggregation-competent conformer to bind it, yet sufficiently altered to preclude proper nucleation or propagation of the amyloid. Morphological analysis by TEM showed shorter and fewer wild-type α-synuclein protofibril assemblies in the presence of **29-71** or **44-54** (**Figure 6e-h**). This supports the hypothesis that these two non-aggerating variants interact with wild-type α-syn oligomeric nuclei or fibril ends, thereby preventing additional wild-type α-syn monomers from assembling onto these structures to create extended fibrils. The kinetic analysis corroborates our findings, indicating that **29-71** substantially inhibits the rate of primary nucleation, while **44-54** specifically suppresses secondary nucleation.

Still more surprising was the behavior of the control variant **64-81**, which formed amyloid fibrils despite the two residues being far apart in the structures of both major polymorphs of α-synuclein. This unexpected observation highlights the complexity of α-synuclein’s conformational landscape. Two recent publications have classified all known α-syn polymorphs according to structural similarity [27, 28]. We examined representative polymorph structures from every cluster in both schemes, which include not only human but even murine α-syn polymorphs. Residues 64 and 81 are far apart in all these structures (**Figure 4**). Thus, variant **64-81** appears to be aggregating in a previously unknown polymorph class. We were able to verify that distinct variants produce morphologically distinct aggregates (**Figure 3b-h**). The observed morphology of **64-81** aggregates was qualitatively distinct from any of the others and, to our knowledge, from any previously reported TEM morphology of α-syn amyloids. This unique morphology lends support to the hypothesis of a new polymorph class. The fact that **64-81** was one of only six variants we tested suggests that many aggregation-competent regions of α-synuclein’s conformational landscape have yet to be mapped. The WT protein may sample them only rarely or under specific *in-vivo* conditions, but the conformational constraint imposed by the disulfide bond may favor and reveal them.

Conformations within aggregate nuclei may differ from those found in mature amyloid fibrils. Moreover, the relative prevalence of distinct polymorphs may evolve during aggregate maturation [48]. Future studies could use disulfide engineering to dissect or control such temporal evolution of conformer or polymorph populations. The physiological effects of distinct disulfide-constrained conformers were also beyond the scope of the present study because disulfide bonds are typically reduced in the cytoplasm. However, disulfide engineering could guide future investigations of cell-level effects of conformational constraints using non-reversible crosslinks in place of disulfides.

## Materials and Methods

### Protein expression and purification

The plasmid T7-7 containing the WT human α-synuclein gene was a kind gift from Prof. Elizabeth Rhoades (U. Penn.). The six double-Cys variants were constructed as follows. The backbone of pET16b plasmid containing a superfolder GFP gene (sGFP) was linearized by PCR with primers designed to omit the sGFP gene. Synthetic gene fragments (Twist) with ends homologous to the linearized pET16b backbone ends were assembled with the linearized backbone by *in vivo* homologous recombination as described previously [49].

The WT protein was expressed and purified as follows. Escherichia coli BL21 (RIL) codon plus cells transformed with T7-7 were grown overnight at 37 °C in TB medium containing 100 µg/ml ampicillin and then subculture into 600 ml of TB containing 100 µg/ml ampicillin. At OD_600_ = 0.7 the cells were induced by adding IPTG at a final concentration 1 mM and kept it overnight at 16 °C. The cells were then palleted down at 4000 rpm and resuspended it into lysis buffer (50 mM tris pH 8.0, 100 mM NaCl, 10 mM EDTA). They were lysed by sonication (40% amplitude with pulse 3s on and 1s off for 10 min) twice. The lysates were then heated at 95 °C for 20 min and centrifuged at 14000 rpm and 4 °C for 30 min. (136 µl/ mL of supernatant) Streptomycin sulfate [10% (w/w)] and glacial acetic acid (228 µl/mL of supernatant) were added to the supernatant. The solutions were then centrifuged at 14000 rpm at 4 °C and the supernatant saved. Equal volume of saturated ammonium sulfate was added to the supernatant and kept it at 4 °C overnight. The solutions were then centrifuged at 4000 rpm and 4 °C for 45 min and the pellets saved. The pellets were resuspended in 100 mM ammonium acetate, and an equal volume of ethanol was added and kept it on ice. Within 1h a white precipitate was observed. The solution was then centrifuged at 4000 rpm at 4 °C for 1h, and the pellet was saved. The pellet was resuspended in 100 mM ammonium acetate buffer and 2M guanidium hydrochloride added to the solution and stored at -80 °C. The protein was finally purified by size exclusion chromatography using 20 mM Phosphate buffer pH 7.0 before use.

Variants **22-33**, **44-54** and **64-81** were expressed and purified exactly as the WT. Variants **46-80**, **69-91**, and **29-71** were expressed with a slightly modified procedure, as follows. The bacterial colonies were plated on Ampicillin plates, and their plasmids were sequenced by GENEWIZ. pET16b plasmids containing the genes of variants were transformed into BL21(DE3) and selected with 100 µg/mL Ampicillin. Single colonies were used to inoculate 10mL TB (Casein digest peptone 12 g/L, Yeast extract 24 g/L, Dipotassium Phosphate 9.5 g/L, Monopotassium Phosphate 2.2 g/L) in 50 mL conical flask which were cultured 6h at 37C, 220 rpm. After 6h cultures were used to inoculate main cultures (1 to 100). Cultures grew at 37 °C and 220 rpm overnight 17-18 h and purified by the above-described protocol.

### Desalting and buffer exchange

To remove ammonium sulfate and exchange the protein into 20 mM PB buffer pH 7.0, 1mM sodium azide, the stored protein stocks were desalted on a HiPrep 26/10 Desalting column (53 mL bed volume, Sephadex G-25 Fine; Cytiva) connected to an ÄKTA GO chromatography system. The column was equilibrated with at least two column volumes of 20 mM PB, pH 7.0, 1 mM sodium azide) at a flow rate of 5–10 mL/min, below the manufacturer’s recommended maximum flow rate (15 mL/min).

For each run, up to 15 mL of samples (≤1/3 of the column bed volume) were applied, in accordance with the recommended sample volume range for HiPrep 26/10 (Cytiva) Desalting columns. Samples were loaded using SEC buffer, and elution was carried out isocratically in the same buffer. UV absorbance at 280 nm was monitored throughout the run. Early eluting fractions corresponding to the void volume peak (containing desalted protein) were collected, whereas later fractions containing small molecules and salts (including ammonium sulfate) were discarded.

### Ion Exchange chromatography

The desalted proteins were purified using QFF 16/10 ion-exchange column (15 ml, HiPrep^TM^, Cytiva). Before sample loading the column was equilibrated using 20 mM Tris-HCl, pH 8.0.15 ml of proteins were loaded followed by washing and then eluted out using a gradient of (0-500 mM NaCl). The eluted protein was concentrated by ultrafiltration (Millipore) and either flash-frozen in liquid nitrogen or directly stored at -80 °C.

### Size Exclusion chromatography

Final purified alpha synuclein variants were achieved by SEC on an ÄKTA Go system equipped with Superdex^TM^ 200 increase 10/300 GL column (Cytiva). Before sample injection, the column was equilibrated with SEC buffer (20 mM Phosphate buffer, 1 mM NaN3, pH 7). Elution was monitored by absorbance at 280 nm, and fractions were collected automatically according to the absorbance profile. Fractions corresponding to the major monomeric peak were analyzed by SDS-PAGE.

### Generating oxidized forms of α-synuclein variants

Purified, fully reduced alpha synuclein mutants were incubated at 4□°C with 1□mM GSSG (an oxidizing agent) overnight. Mutant concentrations were kept below 30□µM to allow intramolecular disulfide bond formation. Oxidized mutants were separated using size exclusion chromatography and further analyzed by a PEG-maleimide gel shift assay [24] and mass spectrometry.

### Mass Spectrometry

The oxidized variants **22–33**, **44–54**, and **64–81** were analyzed by mass spectrometry using a Waters Xevo G2-XS system operated in positive ion mode, with data acquired over an *m/z* range of 110–2000 at a scan rate of 0.2 scans s□¹. The remaining oxidized mutants (**46–80**, **69–91**, and **29–71**) were analyzed by high-resolution mass spectrometry using a Bruker Impact II Q-TOF coupled to an Agilent 1290 LC system.

### Aggregation Assays

The aggregation assay of alpha synuclein mutants was carried out using Spectramax iD3 multi-mode microplate reader at 37 °C in 20 mM phosphate buffer pH 7.0, 100 mM NaCl with 20 µM ThT under high shaking mode. The excitation and emission wavelength set at 440 nm and 485 nm respectively. The sample volume of each well was 100 µl containing a 3mm Teflon bead (Caulys Material Lab). Plates were sealed using Microseal^®^ B Adhesive Sealer (*BIO-RAD*) to avoid evaporation.

### Kinetic Analysis

The kinetic data obtained from the aggregation assays were normalized with respect to minimum and maximum ThT fluorescence intensities. Half-times (t_1/2_) were calculated based on the time when fluorescence intensity reached 50% of its maximum value. All experiments were performed with three repeats and analyzed using the AmyloFit 2.0 platform [29].

### Transmission Electron Microscope

Samples for transmission electron microscopy (TEM) were recovered from aggregation reactions and prepared on carbon films on 3-mm 300-mesh copper grids. 10 µl of sample was drop-cast for this purpose. The sample was then stained with 10 μL of 2% (w/v) Phosphotungstic acid (PTA). Excess liquid was removed, and grids were allowed to dry. The images were recorded by FEI BioTwinG2 transmission electron microscope (120 kV).

**Table 1.** Rate constants from global fits using a multistep secondary nucleation kinetic model.

| **44-54 ($\mu\text{M}$)** | $k_+k_n (M^{-2}h^{-2})$ | $k_+k_2 (M^{-3}h^{-2})$ | $K_M$ |
| --- | --- | --- | --- |
| 0 | $1.26 \times 10^6$ | $3.78 \times 10^{12}$ | $7.17 \times 10^{-11}$ |
| 10 | $8.96 \times 10^5$ | $1.01 \times 10^{12}$ | |
| 20 | $7.31 \times 10^5$ | $9.01 \times 10^{11}$ | |
| 40 | $5.58 \times 10^5$ | $6.22 \times 10^{11}$ | |
| 60 | $6.24 \times 10^5$ | $1.06 \times 10^{12}$ | |
| **29-71 ($\mu\text{M}$)** |  |  |  |
| 0 | $1.26 \times 10^6$ | $3.78 \times 10^{12}$ | $7.17 \times 10^{-11}$ |
| 20 | $6.46 \times 10^5$ | $1.17 \times 10^{12}$ | |
| 40 | $2.58 \times 10^5$ | $3.68 \times 10^{12}$ | |
| 60 | $4.41 \times 10^4$ | $7.16 \times 10^{12}$ | |
| 80 | $1.29 \times 10^4$ | $5.53 \times 10^{12}$ | |
| **22-33 ($\mu\text{M}$)** |  |  |  |
| 0 | $1.26 \times 10^6$ | $3.78 \times 10^{12}$ | $7.17 \times 10^{-11}$ |
| 10 | $1.11 \times 10^6$ | $2.60 \times 10^{12}$ | |
| 20 | $8.14 \times 10^5$ | $2.40 \times 10^{12}$ | |
| 40 | $9.74 \times 10^5$ | $1.06 \times 10^{12}$ | |
| 60 | $1.01 \times 10^6$ | $9.35 \times 10^{11}$ | |

## Acknowledgements

We thank Prof. Elizabeth Rhoades (U. Pennsylvania) for the kind gift of the WT α-synuclein plasmid. We thank Ali Behboodian (Stony Brook U.) for expert technical assistance with molecular cloning for the construction of the 2-Cys variants. We thank Dr. Jeremy Balsbaugh and Dr. Jennifer Liddle at the University of Connecticut Center for Open Research for mass spectrometry analysis of the WT protein and variants **22–33**, **44–54**, and **64–81**. We thank Dr. Beniam Berhane at the Stony Brook University CASDA Mass Spectrometry Core Facility for mass spectrometry analysis of variants **46–80**, **69–91**, and **29–71**. We thank Yunming Hu at the Stony Brook University Central Microscopy and Imaging Center for assistance with transmission electron microscopy. This work was funded by National Institutes of Health grant R00GM141459 to E. S.

## References

[1] Ullman O, Fisher CK, Stultz CM. Explaining the Structural Plasticity of α-Synuclein. Journal of the American Chemical Society. 2011;133:19536–46.

[2] Li A, Rastegar C, Mao X. α-Synuclein Conformational Plasticity: Physiologic States, Pathologic Strains, and Biotechnological Applications. Biomolecules. 2022;12:994.

[3] Uluca B, Viennet T, Petrović D, Shaykhalishahi H, Weirich F, Gönülalan A, et al. DNP-Enhanced MAS NMR: A Tool to Snapshot Conformational Ensembles of α-Synuclein in Different States. Biophysical Journal. 2018;114:1614–23.

[4] Deleersnijder A, Gerard M, Debyser Z, Baekelandt V. The remarkable conformational plasticity of alpha-synuclein: blessing or curse? Trends Mol Med. 2013;19:368–77.

[5] Poewe W, Seppi K, Tanner CM, Halliday GM, Brundin P, Volkmann J, et al. Parkinson disease. Nature Reviews Disease Primers. 2017;3:17013.

[6] Aarsland D, Batzu L, Halliday GM, Geurtsen GJ, Ballard C, Ray Chaudhuri K, et al. Parkinson disease-associated cognitive impairment. Nature Reviews Disease Primers. 2021;7:47.

[7] Iwai A, Masliah E, Yoshimoto M, Ge N, Flanagan L, Rohan de Silva HA, et al. The precursor protein of non-Aβ component of Alzheimer’s disease amyloid is a presynaptic protein of the central nervous system. Neuron. 1995;14:467–75.

[8] Spillantini MG, Schmidt ML, Lee VMY, Trojanowski JQ, Jakes R, Goedert M. α-Synuclein in Lewy bodies. Nature. 1997;388:839–40.

[9] Balestrino R, Schapira AHV. Parkinson disease. European Journal of Neurology. 2020;27:27–42.

[10] Dunker AK, Lawson JD, Brown CJ, Williams RM, Romero P, Oh JS, et al. Intrinsically disordered protein. J Mol Graph Model. 2001;19:26–59.

[11] Teixeira JMC, Liu ZH, Namini A, Li J, Vernon RM, Krzeminski M, et al. IDPConformerGenerator: A Flexible Software Suite for Sampling the Conformational Space of Disordered Protein States. J Phys Chem A. 2022;126:5985–6003.

[12] Zhu J, Li Z, Zheng Z, Zhang B, Zhong B, Bai J, et al. Accurate Generation of Conformational Ensembles for Intrinsically Disordered Proteins with IDPFold. Adv Sci (Weinh). 2025;12:e11636.

[13] Serebryany E, Zhao VY, Park K, Bitran A, Trauger SA, Budnik B, et al. Systematic conformation-to-phenotype mapping via limited deep sequencing of proteins. Molecular Cell. 2023;83:1936–52.e7.

[14] Walker S, Ullman O, Stultz CM. Using Intramolecular Disulfide Bonds in Tau Protein to Deduce Structural Features of Aggregation-resistant Conformations*. Journal of Biological Chemistry. 2012;287:9591–600.

[15] Carija A, Pinheiro F, Pujols J, Brás IC, Lázaro DF, Santambrogio C, et al. Biasing the native α-synuclein conformational ensemble towards compact states abolishes aggregation and neurotoxicity. Redox Biology. 2019;22:101135.

[16] Meade RM, Allen SG, Williams C, Tang TMS, Crump MP, Mason JM. An N-terminal alpha-synuclein fragment binds lipid vesicles to modulate lipid-induced aggregation. Cell Reports Physical Science. 2023;4:101563.

[17] Periquet M, Fulga T, Myllykangas L, Schlossmacher MG, Feany MB. Aggregated α-Synuclein Mediates Dopaminergic Neurotoxicity *In Vivo*. The Journal of Neuroscience. 2007;27:3338–46.

[18] Paslawski W, Mysling S, Thomsen K, Jørgensen TJD, Otzen DE. Co-existence of Two Different α-Synuclein Oligomers with Different Core Structures Determined by Hydrogen/Deuterium Exchange Mass Spectrometry. Angewandte Chemie International Edition. 2014;53:7560–3.

[19] Lorenzen N, Nielsen SB, Yoshimura Y, Vad BS, Andersen CB, Betzer C, et al. How epigallocatechin gallate can inhibit α-synuclein oligomer toxicity in vitro. J Biol Chem. 2014;289:21299–310.

[20] Tuttle MD, Comellas G, Nieuwkoop AJ, Covell DJ, Berthold DA, Kloepper KD, et al. Solid-state NMR structure of a pathogenic fibril of full-length human α-synuclein. Nature Structural & Molecular Biology. 2016;23:409–15.

[21] Li B, Ge P, Murray KA, Sheth P, Zhang M, Nair G, et al. Cryo-EM of full-length α-synuclein reveals fibril polymorphs with a common structural kernel. Nature Communications. 2018;9:3609.

[22] Lautenschläger J, Stephens AD, Fusco G, Ströhl F, Curry N, Zacharopoulou M, et al. C-terminal calcium binding of α-synuclein modulates synaptic vesicle interaction. Nature Communications. 2018;9:712.

[23] Guerrero-Ferreira R, Taylor NMI, Mona D, Ringler P, Lauer ME, Riek R, et al. Cryo-EM structure of alpha-synuclein fibrils. eLife. 2018;7:e36402.

[24] Serebryany E, Yu S, Trauger SA, Budnik B, Shakhnovich EI. Dynamic disulfide exchange in a crystallin protein in the human eye lens promotes cataract-associated aggregation. Journal of Biological Chemistry. 2018;293:17997–8009.

[25] Bisaglia M, Trolio A, Bellanda M, Bergantino E, Bubacco L, Mammi S. Structure and topology of the non-amyloid-beta component fragment of human alpha-synuclein bound to micelles: implications for the aggregation process. Protein Sci. 2006;15:1408–16.

[26] Burré J, Sharma M, Südhof TC. Systematic mutagenesis of α-synuclein reveals distinct sequence requirements for physiological and pathological activities. J Neurosci. 2012;32:15227–42.

[27] Connor JP, Radford SE, Brockwell DJ. Structural and thermodynamic classification of amyloid polymorphs. Structure. 2025;33:1793–804 e3.

[28] Milchberg MH, Warmuth OA, Borcik CG, Dhavale DD, Wright ER, Kotzbauer PT, et al. Alpha-synuclein fibril structures cluster into distinct classes. Biophysical Journal. 2025;124:2571–82.

[29] Meisl G, Kirkegaard JB, Arosio P, Michaels TCT, Vendruscolo M, Dobson CM, et al. Molecular mechanisms of protein aggregation from global fitting of kinetic models. Nature Protocols. 2016;11:252–72.

[30] Spillantini MG, Crowther RA, Jakes R, Hasegawa M, Goedert M. α-Synuclein in filamentous inclusions of Lewy bodies from Parkinson’s disease and dementia with Lewy bodies. Proceedings of the National Academy of Sciences. 1998;95:6469–73.

[31] Villar-Piqué A, Lopes da Fonseca T, Outeiro TF. Structure, function and toxicity of alpha-synuclein: the Bermuda triangle in synucleinopathies. J Neurochem. 2016;139 Suppl 1:240–55.

[32] Mehra S, Ahlawat S, Kumar H, Datta D, Navalkar A, Singh N, et al. α-Synuclein Aggregation Intermediates form Fibril Polymorphs with Distinct Prion-like Properties. Journal of Molecular Biology. 2022;434:167761.

[33] Kim C, Haldiman T, Kang S-G, Hromadkova L, Han ZZ, Chen W, et al. Distinct populations of highly potent TAU seed conformers in rapidly progressing Alzheimer’s disease. Science Translational Medicine.14:eabg0253.

[34] Hromadkova L, Siddiqi MK, Liu H, Safar JG. Populations of Tau Conformers Drive Prion-like Strain Effects in Alzheimer’s Disease and Related Dementias. Cells 2022. p. 2997.

[35] Dhakal S, Robang AS, Bhatt N, Puangmalai N, Fung L, Kayed R, et al. Distinct neurotoxic TDP-43 fibril polymorphs are generated by heterotypic interactions with α-Synuclein. Journal of Biological Chemistry. 2022;298:102498.

[36] De S, Wirthensohn DC, Flagmeier P, Hughes C, Aprile FA, Ruggeri FS, et al. Different soluble aggregates of Aβ42 can give rise to cellular toxicity through different mechanisms. Nat Commun. 2019;10:1541.

[37] Dombkowski AA, Sultana KZ, Craig DB. Protein disulfide engineering. FEBS Letters. 2014;588:206–12.

[38] Taguchi Y, Lu L, Marrero-Winkens C, Otaki H, Nishida N, Schatzl HM. Disulfide-crosslink scanning reveals prion-induced conformational changes and prion strain-specific structures of the pathological prion protein PrP(Sc). J Biol Chem. 2018;293:12730–40.

[39] Serebryany E, Woodard JC, Adkar BV, Shabab M, King JA, Shakhnovich EI. An Internal Disulfide Locks a Misfolded Aggregation-prone Intermediate in Cataract-linked Mutants of Human gammaD-Crystallin. J Biol Chem. 2016;291:19172–83.

[40] Norton-Baker B, Mehrabi P, Kwok AO, Roskamp KW, Rocha MA, Sprague-Piercy MA, et al. Deamidation of the human eye lens protein gammaS-crystallin accelerates oxidative aging. Structure. 2022;30:763–76 e4.

[41] Halverson-Kolkind K, Thorn DC, Tovar-Ramirez M, Shakhnovich E, David L, Lampi K. The Eye Lens Protein, gammaS Crystallin, Undergoes Glutathionylation-Induced Disulfide Bonding Between Cysteines 22 and 26. Biomolecules. 2025;15.

[42] Grad LI, Guest WC, Yanai A, Pokrishevsky E, O’Neill MA, Gibbs E, et al. Intermolecular transmission of superoxide dismutase 1 misfolding in living cells. Proc Natl Acad Sci U S A. 2011;108:16398–403.

[43] Khare SD, Ding F, Dokholyan NV. Folding of Cu, Zn superoxide dismutase and familial amyotrophic lateral sclerosis. J Mol Biol. 2003;334:515–25.

[44] Toichi K, Yamanaka K, Furukawa Y. Disulfide scrambling describes the oligomer formation of superoxide dismutase (SOD1) proteins in the familial form of amyotrophic lateral sclerosis. J Biol Chem. 2013;288:4970–80.

[45] Weismiller HA, Holub TJ, Krzesinski BJ, Margittai M. A thiol-based intramolecular redox switch in four-repeat tau controls fibril assembly and disassembly. J Biol Chem. 2021;297:101021.

[46] Krzesinski BJ, Holub TJ, Gabani ZY, Margittai M. Cellular Uptake of Tau Aggregates Triggers Disulfide Bond Formation in Four-Repeat Tau Monomers. ACS Chem Neurosci. 2025;16:171–80.

[47] Jiang C, Chang J-Y. Isomers of Human α-Synuclein Stabilized by Disulfide Bonds Exhibit Distinct Structural and Aggregative Properties. Biochemistry. 2007;46:602–9.

[48] Wilkinson M, Xu Y, Thacker D, Taylor AIP, Fisher DG, Gallardo RU, et al. Structural evolution of fibril polymorphs during amyloid assembly. Cell. 2023;186:5798–811 e26.

[49] Behboodian A, Serebryany E. Enzyme-free biochemical production of seamlessly N-to-C cyclized peptides from natural or recombinant proteins. Methods Enzymol. 2025;723:427–53.

