## Supporting Information for "Disulfide engineering reveals unexpected pro- and anti-aggregation conformers of human α-synuclein"

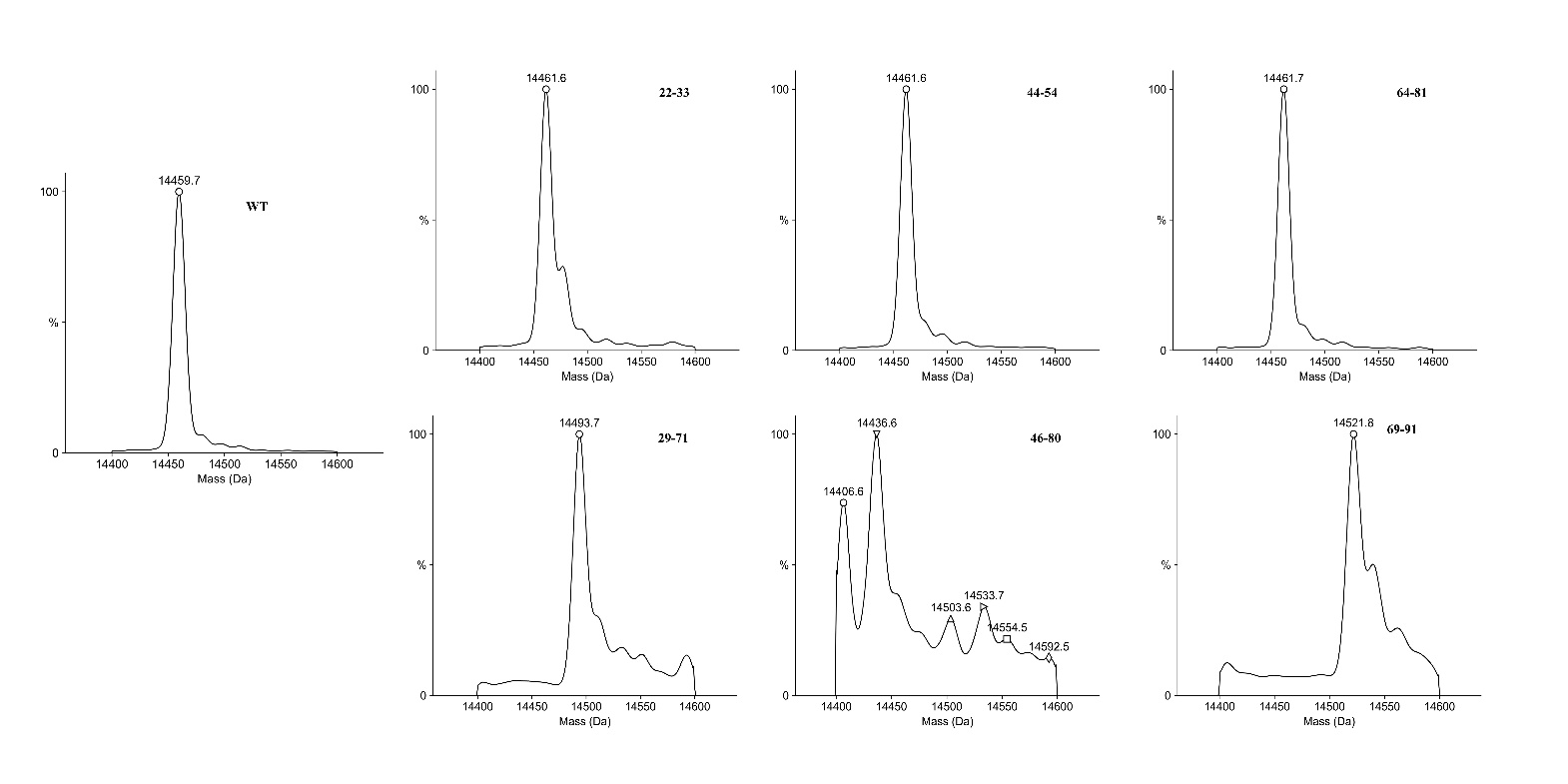

**Figure S1: Deconvoluted electrospray ionization intact high-resolution mass spectra of all protein constructs used in the present study.**

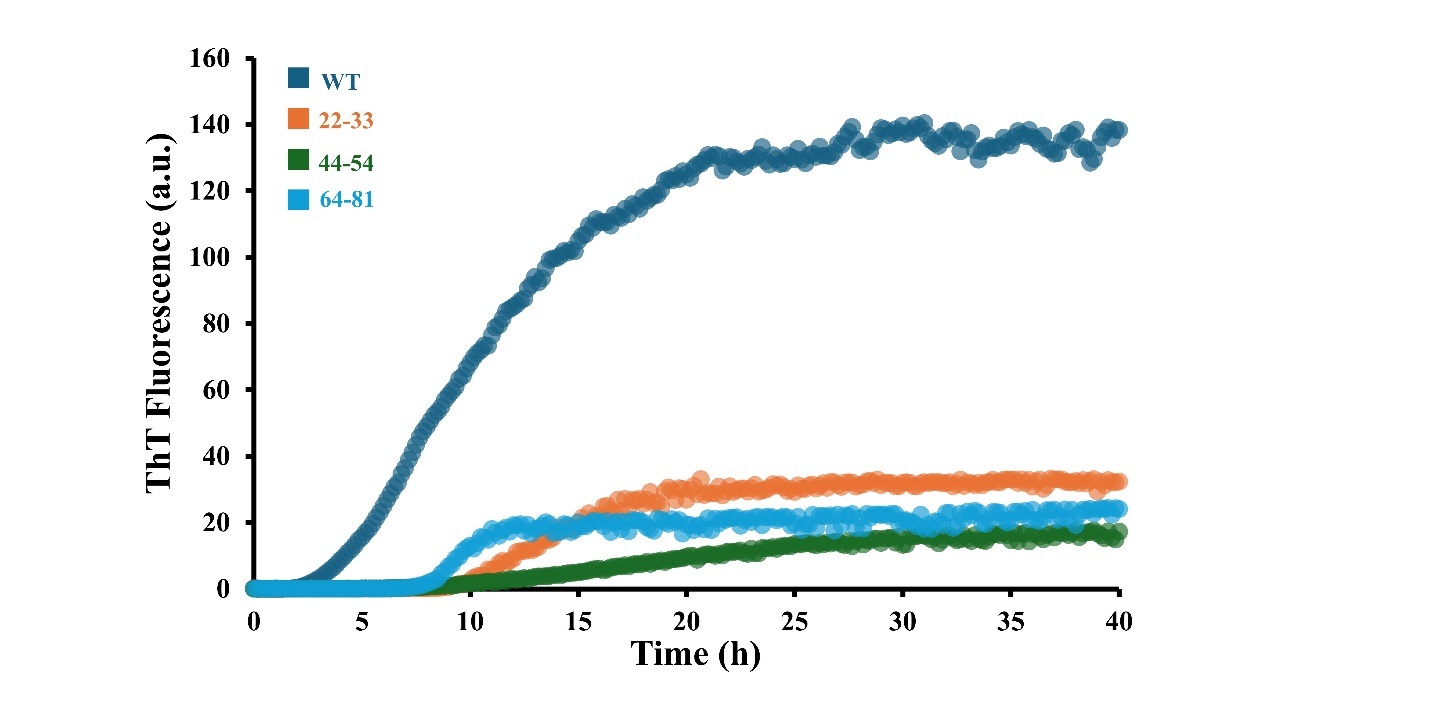

**Figure S2: Amyloid aggregation kinetic traces of variants that did not aggregate when disulfide-crosslinked show aggregation in the presence of reducing agent (TCEP).** Note that all protein concentrations were the same. The variants’ smaller maximum aggregation amplitude relative to the WT could be due to incomplete reduction of the aggregation-preventing internal disulfide bonds. However, we cannot rule out a reduction in aggregation propensity due to the mutations themselves. Most important is the qualitative result that aggregation ability was rescued upon disulfide reduction.

**Table S1: Predicted and observed molecular masses of the α-synuclein constructs**

| Construct | Reduced Molecular Weight (Da) | Oxidized Molecular Weight (Da) | Observed Molecular weight (Da) |
| --- | --- | --- | --- |
| **WT** | 14,460.2 | ------- | 14,459.7 |
| **22-33** | 14,464.2 | 14,462.2 | 14,461.6 |
| **44-54** | 14,464.2 | 14,462.2 | 14,461.6 |
| **64-81** | 14.464.2 | 14,462.2 | 14,461.7 |
| **46-80** | 14,409.1 | 14,407.1 | 14,406.6 |
|  |  |  | 14,436.6* |
| **69-91** | 14,524.3 | 14,522.3 | 14,521.8 |
| **29-71** | 14,496.2 | 14,494.2 | 14,493.7 |

*Two major peaks were observed in the deconvoluted mass spectrum of this variant (Fig. S2). However, sample desalting in this particular analysis appears to have been poor, since mass shifts consistent with phosphate or sulfate adducts were also found. Thus, the +30 Da. shift could be due to a chemical contaminant from prior mass spectrometry runs, e.g., formaldehyde.
